# Long-range order from local interactions: organization and development of distributed cortical networks

**DOI:** 10.1101/335828

**Authors:** Gordon B. Smith, Bettina Hein, David E. Whitney, David Fitzpatrick, Matthias Kaschube

## Abstract

The cortical networks that underlie behavior exhibit an orderly functional organization at local and global scales, which is readily evident in the visual cortex of carnivores and primates^1-6^. Here, neighboring columns of neurons represent the full range of stimulus orientations and contribute to distributed networks spanning several millimeters^2,7-11^. However, the principles governing functional interactions that bridge this fine-scale functional architecture and distant network elements are unclear, and the emergence of these network interactions during development remains unexplored. Here, by using in vivo wide-field and 2-photon calcium imaging of spontaneous activity patterns in mature ferret visual cortex, we find widespread and specific modular correlation patterns that accurately predict the local structure of visually-evoked orientation columns from the spontaneous activity of neurons that lie several millimeters away. The large-scale networks revealed by correlated spontaneous activity show abrupt ‘fractures’ in continuity that are in tight register with evoked orientation pinwheels. Chronic in vivo imaging demonstrates that these large-scale modular correlation patterns and fractures are already present at early stages of cortical development and predictive of the mature network structure. Silencing feed-forward drive through either retinal or thalamic blockade does not affect network structure suggesting a cortical origin for this large-scale correlated activity, despite the immaturity of long-range horizontal network connections in the early cortex. Using a circuit model containing only local connections, we demonstrate that such a circuit is sufficient to generate large-scale correlated activity, while also producing correlated networks showing strong fractures, a reduced dimensionality, and an elongated local correlation structure, all in close agreement with our empirical data. These results demonstrate the precise local and global organization of cortical networks revealed through correlated spontaneous activity and suggest that local connections in early cortical circuits may generate structured long-range network correlations that underlie the subsequent formation of visually-evoked distributed functional networks.

---

The rules that govern how neurons in cortical columns participate in the larger distributed networks that they comprise remain poorly understood. Anatomical studies that have probed the organization of horizontal connections in visual cortex suggest that network interactions could exhibit considerable functional specificity^9-11^. But the fine scale structure of network interactions, and the degree to which the activity of a given cortical locus is reliably coupled with the spatiotemporal patterns of activity elsewhere in the network, have yet to be determined. Likewise, the sequence of events that leads to the development of mature network interactions is largely unexplored since these occur at early stages in development when visual stimuli are ineffective in evoking reliable neuronal responses^12,13^. Previous studies have suggested that spontaneous activity patterns may be a powerful tool for probing network structure independent of stimulus-imposed organization, and one that is applicable especially early in development^14-18^. This approach is further supported by the finding that under anesthesia individual spontaneous events can resemble visually-evoked activity patterns for stimuli known to engage distributed functional networks^19,20^.

We therefore sought to exploit the sensitivity and resolution of *in vivo* calcium imaging of ongoing spontaneous activity in the ferret visual cortex to gain new insights into fundamental questions of network interactions and their formation over development. In the awake visual cortex imaged near the time of eye-opening, wide-field epifluorescence imaging reveals a highly dynamic modular pattern of spontaneous activity that covers millimeters of cortical surface area (Fig. 1a,b; Extended Data Movie 1). Spontaneous activity patterns consist of a distributed set of active domains which become active either near simultaneously or in a spatial-temporal sequence spreading across the field of view within a few hundred milliseconds.

**Figure 1:**
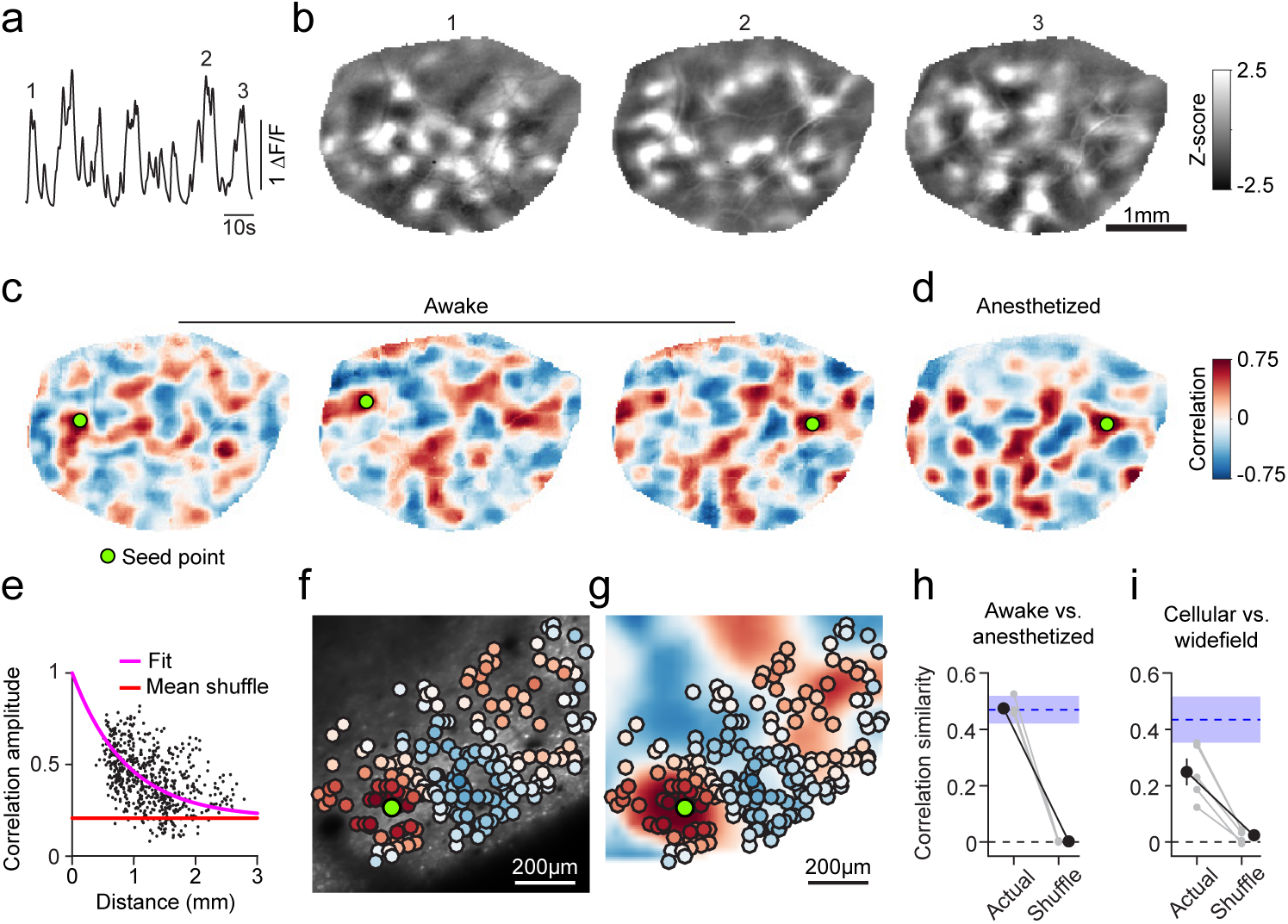
Correlated spontaneous activity in awake ferret visual cortex reveals large-scale modular distributed functional networks. **a.** Timecourse of spontaneous activity measured with wide-field epifluorescence in an awake ferret (mean across pixels in ROI). **b**. Representative z-scored images of spontaneous events at times indicated in (**a**). **c**. Spontaneous activity correlation patterns shown for 3 different seed points (green circle). Correlation patterns span millimetres, can show both rapid changes between nearby seed points (*left* and *middle*) and long-range similarity for distant seed points (*middle* and *right*). **d**. Correlation patterns are highly similar in the awake and anesthetized cortex. **e**. Correlation values at maxima as a function of distance from the seed point showing that correlation amplitude remains strong over long distances. **f-g**. Spontaneous activity is modular and correlated at the cellular level (**f**) and shows good correspondence to spontaneous correlations obtained with wide-field imaging (**g**). **h**. Correlations measured under anaesthesia are statistically similar to those in the awake cortex (n=5, grey: individual animals, black: mean ± SEM). Blue shaded region indicates within-state similarity (mean ± SEM). **i**. Cellular correlations are significantly similar to wide-field correlations (n=5, grey: individual animals, black: mean ± SEM). Blue region indicates within-modality similarity (mean ± SEM).

The strikingly regular modular structure of the spontaneous activity patterns suggests a high degree of correlation in the activity of specific subsets of neurons that are part of this distributed network. To evaluate this correlation structure, we first detected individual large-scale spontaneous events within ongoing spontaneous activity (see Methods), which occurred frequently in the awake cortex (inter-event interval: 2.13 (1.33 – 6.53) seconds; duration 1.13 (0.73 – 1.73) seconds; median and IQR; n=5 animals, Extended Data Fig. 1a,b). The spatial structure of activity was relatively stable with minor fluctuations over the course of an event, and exhibited frame-to-frame cross-correlations near 0.5 for a two-second window centered on the peak activity (Extended Data Fig. 2). The frequency and duration of spontaneous events is reminiscent of synchronous states observed in LFP recordings from awake animals, appearing distinct from both the desynchronized activity often observed during active attention, as well as the oscillatory activity seen in slow-wave sleep and with certain types of anesthesia^21^.

**Figure 2:**
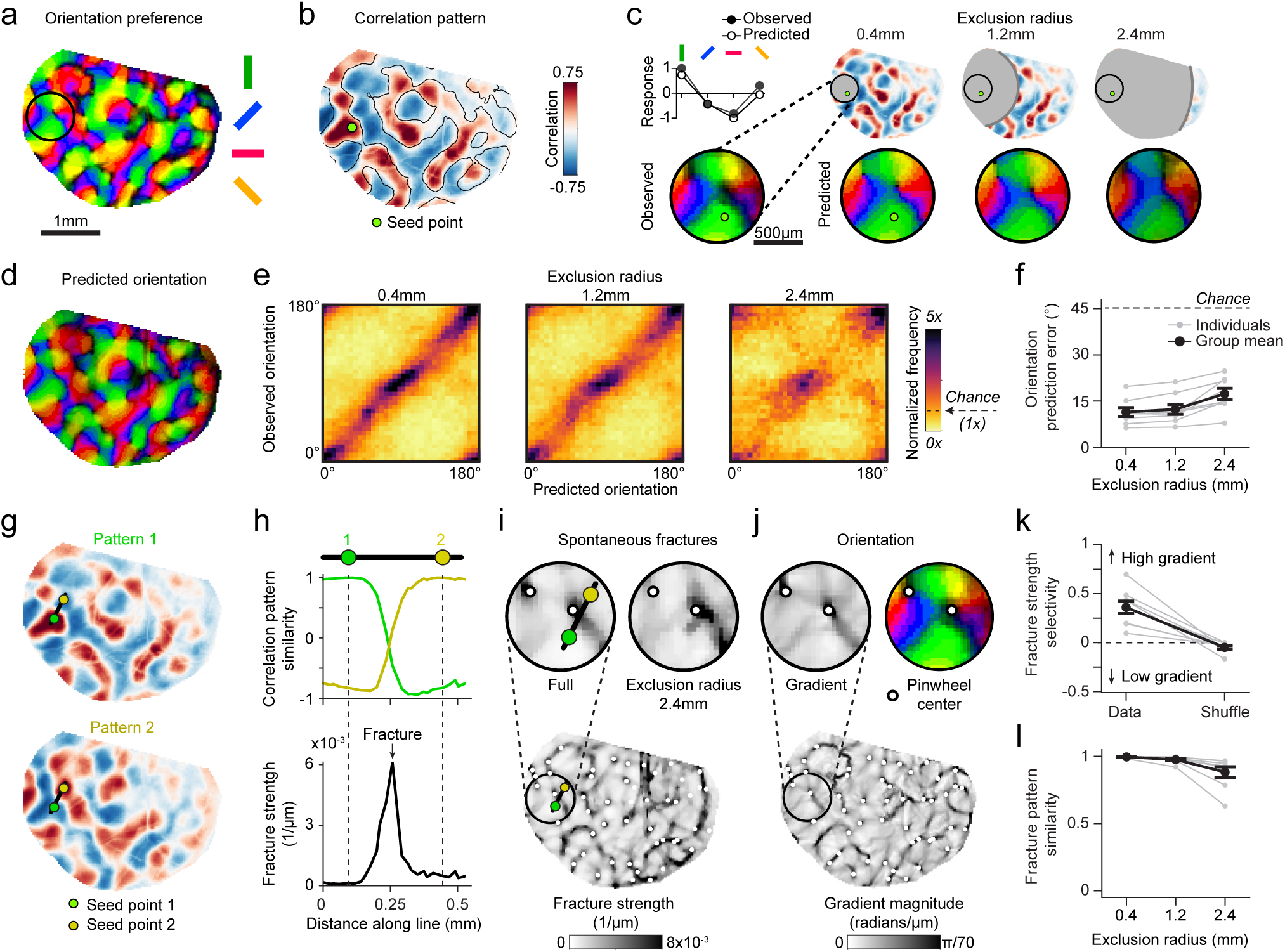
Tight interrelation between long-range spontaneous correlation and the fine-scale structure of orientation columns after eye opening (EO). **a.**Orientation preference map. **b.** Spontaneous correlation pattern for indicated seed point. Contour lines from vertical selective domains from (**a**) reveal that spontaneous correlations closely resemble the layout of orientation preference map. **c**. Local orientation tuning for region within black circle in (**a**) can be accurately predicted from the aggregate orientation tuning of distant cortical locations, weighted by long-range correlations. (*Top left*) Observed and predicted tuning for single pixel shown below. (*Bottom left*) Observed orientation tuning. (*Right*) Accurate orientation predictions based on increasingly distant regions of spontaneous correlations (excluding pixels within either 0.4, 1.2, or 2.4 mm from the seed point). **d.** The prediction based on correlations >1.2 mm away (excluding all correlations <1.2 mm from seed point) matches the actual preferred orientation within the entire field of view (see **(a)**). **e.** Across animals, the precision of predicted orientation tuning remains high, even when based on restricted regions more than 2.4mm away from the site of prediction (see (**c**)) **f**. Prediction error as function of exclusion radius (45° is chance level). **g-h.** Fractures in correlated networks. Advancing the seed-point along the black line in (**g**) reveals a punctuated rapid transition in global correlation structure expressed by a high rate of change in the correlation pattern between adjacent pixels (**h**, bottom). **i**. Locations with high rate of change form a set of lines across the cortical surface, which we termed spontaneous fractures. **j**. The layout of spontaneous fractures precisely coincides with the high-rate of change regions in the orientation preference map. **k.** Correlation fractures show selectivity for regions of high orientation gradient. **l.** Fracture location is independent of local correlation structure and remains stable when only long-range correlations are included (see **c**). For e, f, k, l: n = 8 animal experiments with 5 days or more of visual experience (gray); group data in f,k,l is shown as mean ± SEM (black).

Spontaneous activity correlation patterns were then computed from detected events by choosing a given seed point and computing its correlation in spontaneous activity with the remaining locations within the field of view. Correlation patterns for a given seed point show a striking widespread modular organization, with patches of positively correlated activity separated by patches of negatively correlated activity (Fig. 1c). Correlation patterns exhibited significant long-range structure, with statistically significant correlations persisting for more than 2 mm (Fig. 1e; p<0.01 vs. surrogate for example shown, p<0.01 for 10 of 10 animals imaged following eye-opening). The consistency of the correlation patterns is evident in the fact that nearby seed points placed in regions that are negatively correlated exhibit dramatically different spatial correlation patterns (Fig. 1c, *left* and *middle*), while seed points placed millimeters away in regions that are positively correlated show quite similar spatial correlation patterns (Fig. 1c, *middle* and *right*). Moving the seed point across the cortical surface revealed a large diversity of correlation patterns (Extended Data Movie 2), consistent with principal component analysis revealing that the overall number of relevant global variance components in spontaneous activity patterns is typically larger than ten (Extended Data Fig. 3; 13±3 PCs required to explain 75% variance, mean ± standard deviation, n=10).

**Figure 3:**
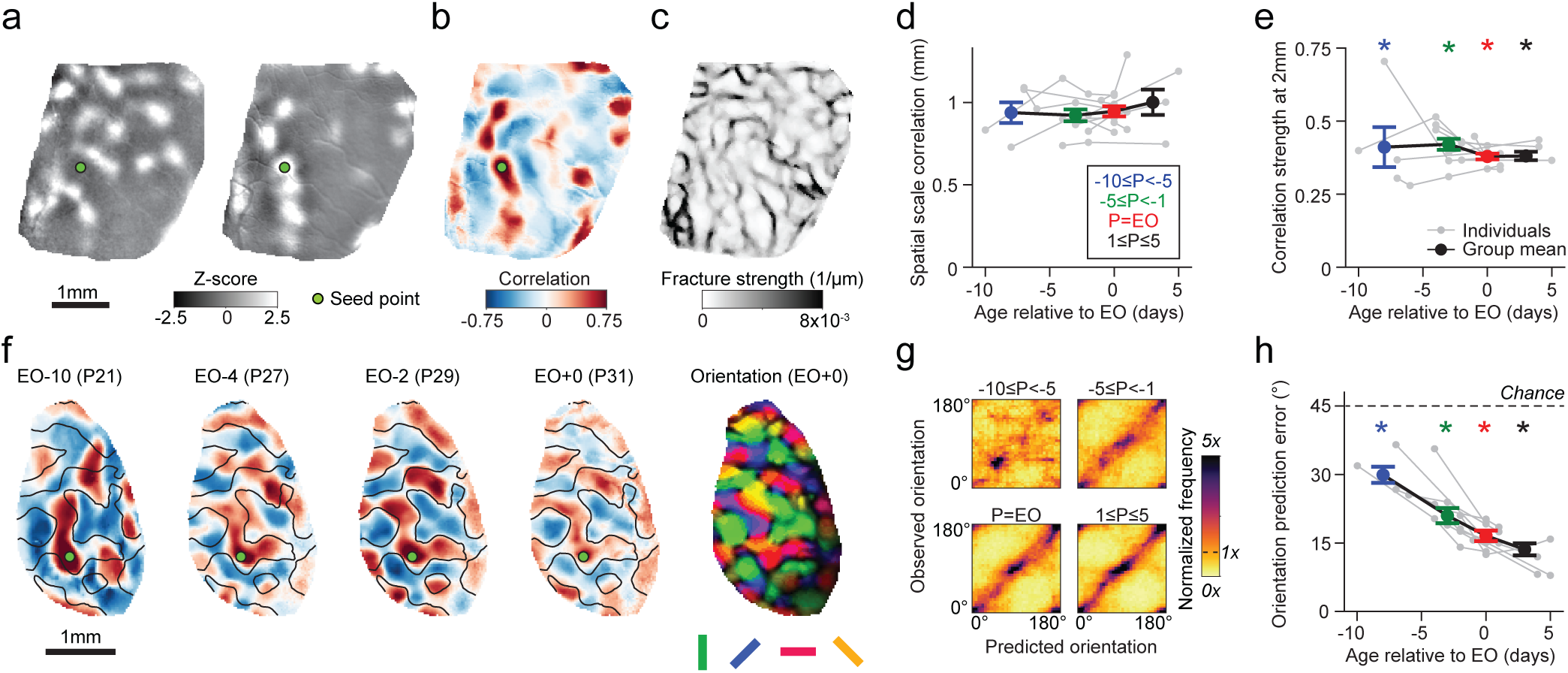
Early spontaneous activity exhibits long-range correlations and predicts future evoked responses. a. Representative z-scored images of early spontaneous activity at P23, seven days prior to EO. **b-c.** Early spontaneous activity shows hallmarks of mature spontaneous activity, including long-range correlated activity **(b)** and pronounced spontaneous fractures (**c**). **d.** The spatial scale of correlations in spontaneous activity (decay constant of correlation maxima) does not change across ages. P denotes age relative to EO. **e.** The magnitude of long-range correlations for maxima 2mm from the seed point is statistically significant at all ages examined. **f.** Longitudinal imaging of a chronically-implanted animal reveals that early spontaneous correlation patterns exhibit signatures of the mature orientation map (*right*), despite considerable reorganization in correlation structure. Contour lines indicate horizontal selective domains measured at EO. **g.** The structure of spontaneous correlations can predict the future mature orientation preference map organization as early as 10 days before eye opening. P denotes age relative to EO. **h.** Spontaneous correlation structure predicts orientation preference significantly better than chance, even at the youngest ages examined. For d, e: n=10 and g, h: n=11 chronically recorded animals; e, h: asterisks indicate p<0.0001, actual vs surrogate data; d,e,h: group data is shown as mean ± SEM.

To determine the impact of brain state and anesthesia on the spatial patterns of correlated spontaneous activity, we followed awake imaging sessions with imaging under light anesthesia (0.5-1% isoflurane). Notably, although spontaneous activity in the awake cortex was more dynamic than under anesthesia, (Extended Data Fig. 4a,c; Extended Data Movie 3), the spatial patterns of spontaneous activity, both in extent, modularity, and correlation structure were remarkably similar across states (Fig. 1d,h; Extended Data Fig. 4; p=0.031 one-sided Wilcoxon signed-rank test, with 5 of 5 experiments from 3 animals individually significant at p<0.001 vs. shuffle). Given this strong similarity, awake and anesthetized recordings were pooled in subsequent analyses, and anesthetized recordings were performed exclusively in some experiments.

**Figure 4:**
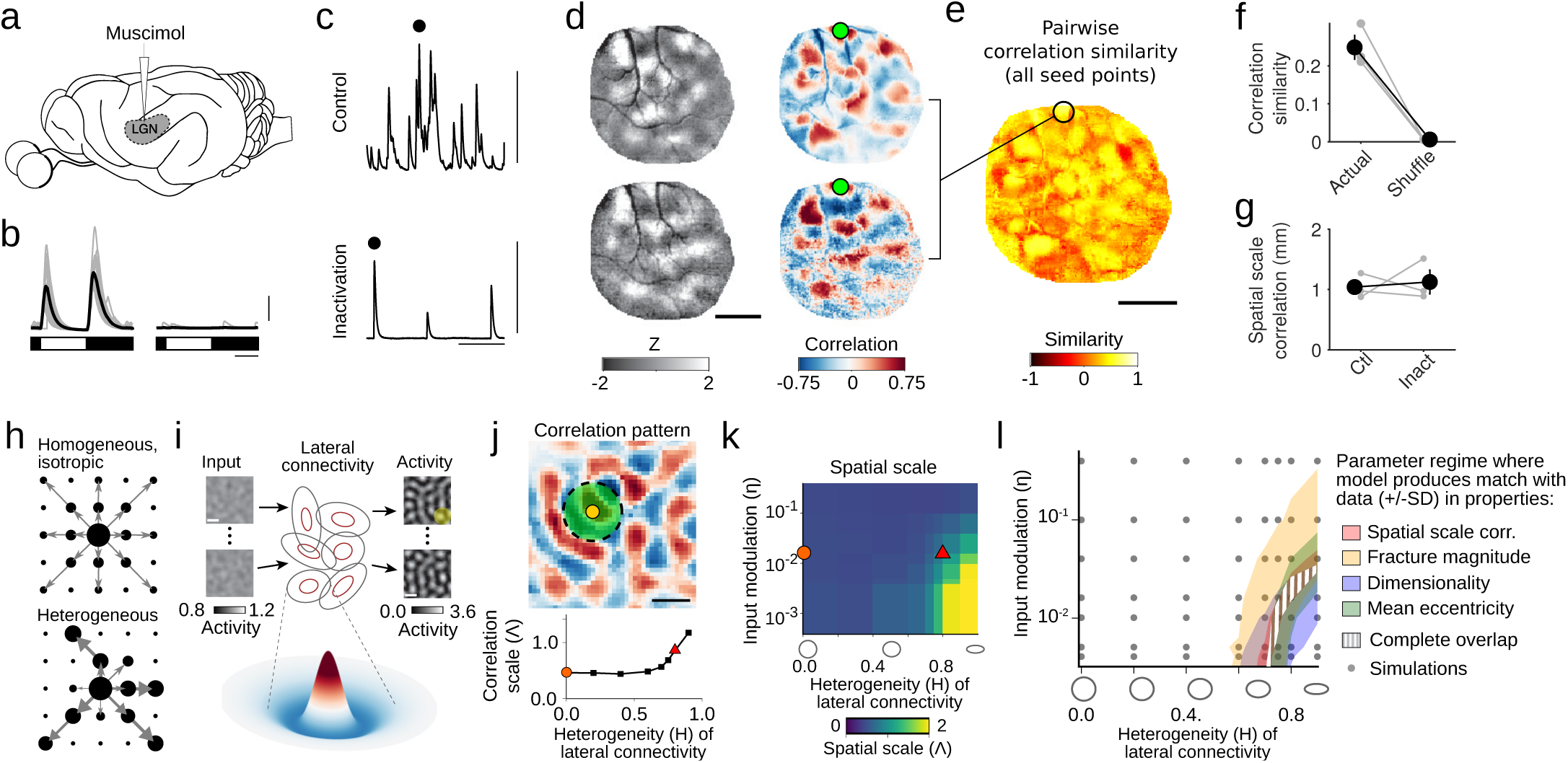
Circuit mechanisms for long-range correlations in early visual cortex. a-g. Long-range correlations in spontaneous activity persist in the absence of feed-forward input. **a**. Cortical spontaneous activity was measured before and following LGN inactivation via targeted muscimol infusion. **b**. Cortical responses (averaged across all pixels in ROI) to full-field luminance changes before (*left*) and after (*right*) LGN inactivation. Scale bars: 5 sec, 0.5 δF/F. **c**. Time-course of spontaneous activity for mean of all pixels before (*top*) and after (*bottom*) inactivation. Scale bars: 30 sec, 0.5 δF/F. **d**. Representative spontaneous events (*left*) and correlation patterns (*right*) before (*top*) and after (*bottom*) inactivation. **e**. Similarity of correlation structure in representative experiment before and after inactivation for all cortical locations. **f**. Correlation structure was significantly more similar before and after inactivation than shuffled data (p<0.01, for 3 of 3 individual experiments, bootstrap test). **g**. The spatial scale of spontaneous correlations remains long-range following LGN inactivation. **h.** Homogenous local connections (arrow) induce moderate correlations with all nearby domains (black dots), whereas heterogeneity introduces biases, favouring some correlations (large dots) more than others (small dots). **i**. A dynamical circuit model of spontaneous activity in the early cortex: a constant input modulated spatially by filtered noise is fed into a recurrent network with short-range, heterogeneous Mexican-hat (MH) connectivity. It produces a set of modular output patterns with typical spatial scale Λ determined by the MH size (average MH size illustrated by the yellow circle). **j.** Correlation pattern in model obtained for heterogeneity *H*=0.8 and input modulation η=0.016 (SD of noise component) shows long-range correlations in agreement with experiment (n=100 output patterns, 16% of modelled region shown) (*top*). The spatial scale of correlations increases with increasing heterogeneity in the lateral connections (*bottom,* η=0.016). **k.** The scale of correlations also increases with decreasing input modulation. Red triangle in (**j,k**): parameters for correlation pattern shown in (**j**, top). Orange circle in (**j,k**): isotropic, homogeneous connectivity, inconsistent with the correlation scale in experiment (compare **j**, bottom, and Fig. 3d). **l.** In the parameter regime where the model spontaneous patterns approach the empirically observed dimensionality, their short- and long-range correlation structure is in quantitative agreement with the experimental data. Shaded regions show parameter regimes in the model in which different properties lie within the range (mean ± SD) of the experimental values (using 1Λ=1mm, linear interpolation between simulations). Scale bars: domain spacing 1Λ (**i**,**j**,**l**); 1mm (**d**,**e**).

We next performed 2-photon imaging with cellular resolution in conjunction with wide-field imaging in the same animal, finding strong and spatially organized spontaneous activity at the cellular level. The duration of events was similar to that observed with wide-field imaging (0.88 (0.54–1.32) seconds, median and IQR), and within an event the pattern of active cells was largely consistent across time (frame-to-frame correlations >0.5 for one second around the peak frame within an event, p<0.01 vs. random epochs, bootstrap test). Cellular spontaneous events exhibited similar durations to events detected in wide-field data (Extended Data Fig. 1; Extended Data Movie 4). The modular organization of spontaneous activity and the spatial correlation patterns observed in populations of individual layer 2/3 neurons was well-matched to those found with wide-field imaging, demonstrating that the network structures revealed with wide-field epifluorescence imaging reflect the spatial activity patterns of individual neurons in superficial cortex (Fig. 1f-g, i; p=0.031 one-sided Wilcoxon signed-rank test, with 4 of 5 experiments from 3 animals individually significant at p<0.05 vs. shuffle). Taken together, these results indicate that neurons in layer 2/3 of visual cortex participate in long-range modular networks whose correlation structure is determined by circuit elements that are independent of brain state.

As individual spontaneous events can resemble the patterns of activity evoked by oriented stimuli^19,20^, we sought to determine whether this correlated spontaneous activity, representing an average over many events and therefore potentially revealing the underlying network architecture, accurately reflects the fine-scale structure of the visually evoked modular network that represents stimulus orientation. We first compared the patterns of spontaneous correlations to the spatial layout of visually-evoked orientation domains in animals imaged 5 or more days after eye-opening, when orientation selectivity is robust (Fig. 2a). We observed seed points for which the spontaneous correlation pattern closely matched the layout of orientation domains, even at remote distances from the seed point (Fig. 2b, Extended Data Fig 5, mean similarity of orientation vs. spontaneous: |r | = 0.42 ± 0.03; mean ± SEM; n=8). We also found a significantly weaker but above chance similarity of spontaneous correlations to the ocular dominance map (Extended Data Fig 5, mean similarity of OD vs. spontaneous: |r | = 0.18 ± 0.04; mean ± SEM; n=3; p<0.0001 vs. surrogate for 3 of 3 animals tested; p=0.02, Mann-Whitney). The strong long-range similarity to orientation preference for certain seed points suggests that the orientation tuning at such seed points can be predicted from the tuning at remote locations that are correlated in spontaneous activity. To test this idea, we computed the sum over tuning curves at distant locations weighted by their spontaneous correlation with the seed point and compared this prediction to the seed point’s actual tuning curve (Fig. 2c, *top left*). Correlated spontaneous activity predicted the preferred orientation in a small circular patch of radius 0.4mm with a high level of accuracy. Notably, orientation predictions remained highly accurate even when only considering correlations in regions more than 2.4mm away from the circle’s center point (Fig. 2c-f, p<0.0001 vs. surrogate for all exclusion radii, with 8 of 8 individual animals significant at p<0.05 across all exclusion radii), demonstrating a high degree of long-range fidelity in the structure of spontaneously active networks and those evoked through oriented visual stimuli.

Orientation preference displays pronounced heterogeneity in rate of change across the cortical surface, most notably at pinwheel centers^2,7,22^; thus an even more stringent test of the relation of the spontaneous activity to the fine structure of the orientation map is to ask whether spontaneous correlation patterns exhibit an analogous heterogeneity in their rate of change that correlates with the orientation preference map. Moving the seed point across the cortex shows regions of gradual change in correlation structure punctuated by abrupt shifts in the large-scale pattern (Extended Data Movie 2). By computing the rate of change of the correlation pattern as the seed point was moved (see Methods), we observed peaks of large change over relative small distances (Fig. 2g-h). A systematic mapping across the cortical surface revealed a set of lines, which we termed spontaneous fractures (Fig. 2i). Moving the seed point across any of these fractures led to strong changes in the global correlation pattern, while correlations changed relatively little when the seed point was moved within the regions between the fractures. Notably, the layout of spontaneous fractures was stable even when only correlations with remote locations (>2.4 mm from seed point) were used to predict the local rate of change (Fig. 2i,l; correlation between fracture patterns for full area vs. >2.4mm: r=0.88±0.04, mean ± SEM, n=8). The layout of spontaneous fractures followed closely the heterogeneity in the rate of change in preferred orientation (Fig. 2j), which also formed an intricate network of lines across the cortical surface and often appeared in tight register with one another (Fig. 2k; p = 0.0078, Wilcoxon signed-rank test, with 8 of 8 individual animals significant at p<0.001 vs. shuffle), as highlighted by the positions of pinwheel centers (Fig. 2i,j). Thus spontaneous fractures are local manifestations of dramatic large-scale diversity in distributed network structure and emphasize that both the fine- and large-scale organization of correlated spontaneous activity are closely matched with the structure of the visually-evoked orientation network.

Having established that the spontaneous correlation structure faithfully captures key aspects of the distributed networks evoked by visual stimulation, we sought to exploit the correlation structure to gain insights into the nature of these networks at earlier stages of development and determine how they evolve to the mature state. Surprisingly, even at post-natal day 21, 10 days prior to eye opening (and the earliest time point examined), we observed robust spontaneous activity, which exhibited modular patterns that extended for long distances across the cortical surface (Fig. 3a; Extended Data Movie 5) and with a temporal structure similar to, albeit slightly more static than in older animals (Extended Data Fig. 2c-e). Likewise, we found strong correlation patterns that displayed pronounced peaks even several millimeters away from the seed point (Fig. 3b), consistent with electrophysiological recordings^23^. Indeed, the spatial scale of spontaneous correlations changed very little with age and already 10 days prior to eye opening it was nearly as large as 5 days after eye opening (Fig. 3d,e; correlation spatial scale: p = 0.86, Kruskal Wallis H-test; correlation strength at 2 mm: p<0.0001 vs. surrogate for all groups; across groups: p = 0.42, Kruskal Wallis H-test). Moreover, spontaneous fractures were already pronounced at the earliest time points, indicating the presence of locally highly organized long-range functional networks in the early cortex (Fig. 3c). These observations are surprising in light of the limited development of long-range horizontal connections at this early age. Anatomical studies in ferret visual cortex show that layer 2/3 pyramidal cell axons exhibit only about two branch points at P22^24^, extend only up to 1mm from the cell body^25^, and are still lacking spatial clusters of synaptic terminals, which are distributed across several millimeters in the mature cortex^26^, but only start to become evident at about P26-27^25,27^.

Our finding that modular activity, long-range correlations, and fractures—all the features that define the modular distributed network—are evident at this early age could suggest that the basic structure of the network may already be similar to its mature state. If so, then we should be able to predict the structure of the mature visually evoked network from the spontaneous activity correlation patterns at these early time points. To test this possibility, we employed chronic recordings starting as early as postnatal day 21, and 10 days prior to eye opening, and mapped all imaging data from each animal onto a common reference frame via an affine transformation, allowing us to track the structure of spontaneous correlations across development (Fig. 3f). We assessed the ability to predict local tuning from remote correlated locations—similar to Fig. 2f, but now applied across age—to predict the visually evoked orientation map from early spontaneous activity. We found that predictions remained fairly accurate up to 5 days prior to eye opening and were above chance even for the youngest age group, showing that even at this early stage, signatures of the future visually-evoked network are apparent (Fig. 3g,h; EO -10 to -5: p<0.0001 vs. surrogate, 4 of 5 individual data points significant vs. surrogate at p<0.05). At the same time, it is clear that there is extensive refinement of the distributed network over this time period (Fig. 3f; Extended Data Fig. 6), such that the ability of the spontaneous correlation patterns to predict the visually-evoked orientation patterns increases significantly with age (Fig. 3h; p=0.0004, Kruskal Wallis H-test; EO -10 to -5 vs. EO: p=0.004, Wilcoxon rank-sum). It is also clear that the refinement during this period involves a rearrangement in the spatial organization of spontaneous fractures (Extended Data Fig. 6c, f).

Having demonstrated that large-scale modular networks are present early in cortical development prior to the maturation of horizontal connectivity, and are linked to the system of orientation columns in the mature cortex, we next considered the potential circuit mechanisms capable of generating such large-scale distributed networks in the early cortex. Spontaneous retinal waves are a prominent feature of the developing nervous system^28^, which exhibit highly organized structure and have been shown to propagate into the visual cortex^29^. To assess a potential contribution of retinal waves to activity pattens in the early cortex, we performed intraocular infusions of TTX, in conjunction with wide-field imaging of spontaneous activity in the cortex. Despite completely abolishing light-evoked responses (response amplitude, pre: 0.357 ± 0.061 ΔF/F, post: 0.023 ± 0.028 ΔF/F, mean ± SEM, bootstrap test vs. baseline: pre-inactivation: p=0.009, post: p=0.365, n=3, P22-25), we continued to observe large-scale spontaneous events, and the spatial correlation structure was significantly more similar to the pre-inactivation structure than would be expected by chance (Extended Data Fig. 7, similarity vs. shuffle, p<0.01 for 3 of 3 animals, bootstrap test).

To address the possibility that coordinated thalamic activity drives large-scale correlations in the early cortex^30^, we infused muscimol into the LGN to silence feed-forward inputs to the cortex at P22-25 (Fig. 4a). Muscimol completely blocked light-evoked responses (Fig. 4b, response amplitude, pre: 0.720 ± 0.105, post: 0.005 ± 0.006, mean ± SEM, bootstrap test vs. baseline: pre-inactivation: p=0.009, post: p=0.258, n=3), and dramatically decreased the frequency of spontaneous events in the cortex (Fig. 4c, < 1 event / minute, with a 713 ± 82 % increase in the inter-event-interval, mean ± SEM, n=3). However, the events remaining after geniculate inactivation still showed large-scale modular activity patterns spanning millimeters, and exhibited spatial correlation structures similar to those observed prior to inactivation (Fig. 4d-f, similarity vs. shuffle: p<0.01 for 3 of 3 individual experiments, bootstrap test), consistent with prior experiments where silencing was induced via optic nerve transection^23^. In addition, we find that the spatial layout of correlation fractures is also similar following LGN inactivation (fracture similarity: 0.164 ± 0.015, p=0.04, bootstrap test, n=3 animals), suggesting that the fine-scale structure of correlation patterns is also generated within cortical circuits. Notably, the spatial extent of correlations was unchanged following muscimol (Fig. 4g, control: 1.04 ± 0.12; inactivation: 1.13 ± 0.20 mm, mean ± SEM), demonstrating that feedforward drive cannot account for the spatial structure and extent of correlated spontaneous activity in the early cortex. These results suggest that rather than being inherited from feed-forward pathways, the modular, large-scale distributed networks present in the early visual cortex are intrinsically generated within cortical circuits.

However, as these large-scale cortical networks are present prior to the maturation and elaboration of long-range horizontal connectivity, these results also present a conundrum. To explore how a developing cortex lacking long-range connectivity could generate long-range correlated patterns of activity, we examined dynamical network models of firing rate units^31^, variants of which have been used previously to model spontaneous activity in the mature visual cortex^32-34^. In these models, modular patterns of activity arise via lateral suppression and local facilitation. Such an interaction is commonly assumed to result from lateral connections that are 1) identical at each position in cortex, 2) circularly symmetric, and 3) follow a ‘Mexican-hat’ profile. Although there is some evidence for Mexican-hat structures in early ferret visual cortex^35^, an anatomical Mexican hat is not strictly required for the formation of modular patterns, which can arise even if the range of lateral excitation exceeds that of inhibition (Extended Data Fig. 8, Supplementary Notes)—an arrangement consistent with recordings in mature cortical slices^36^.

Notably, despite producing modular patterns of activity, the resulting patterns produced by both models exhibit an unrealistic regular hexagonal structure (Extended Data Fig. 8). Additionally, correlation patterns based on this activity are nearly identical across seed points and decay more rapidly with distance, failing to show long-range structure (Extended Data Fig. 8) and correlation fractures, indicating that this mechanism alone cannot account for the widespread and diverse correlation patterns we observe *in vivo*. Rather, the fact that different seed points display different correlation patterns, suggests that local network interactions are not homogeneous across cortex. Instead, if local connections vary, this can bias the interactions between neighboring domains, such that some show a stronger tendency to be co-active than others. Such biases can propagate across the network via multi-synaptic connections and induce correlations even between remote locations (Fig. 4h). Thus, local, but *heterogeneous* synaptic connections may ‘channel’ the spread of activity across cortex, thereby constraining the overall layout of activity patterns, and explaining the observed pronounced correlations found between remote network elements.

To test the idea that heterogeneous local connections can produce long-range correlations, we modeled cortical spontaneous activity using a dynamical rate network^31-34,37^, in which model units (representing a local pool of neurons) receive recurrent input from neighboring units weighted by an anisotropic Mexican-hat function, whose longer axis varies randomly across the cortical surface (Fig. 4i; Methods). To this network, we supplied a constant drive, modulated spatially by a Gaussian random field with only local correlations (Fig. 4i, *left*). For sufficiently strong connections, the network activity evolves towards a modular pattern with roughly alternating patches of active and non-active domains (Fig. 4i, *right*). In the regime of considerable heterogeneous connectivity and moderate input modulation, the model produces pronounced long-range correlations (Fig. 4j,k; Extended Data Fig. 9a,b) and fractures (Extended Data Fig. 9c,d) in quantitative agreement with experiment (Fig. 4l). In contrast, these properties do not match with experiment if lateral connections are homogeneous and isotropic (Fig. 4k, l, left region in diagram). While the dimensionality of the input patterns to the model is high, the dimensionality of the produced set of activity patterns in the heterogeneous regime is much smaller approaching values close to those observed in experiment (Extended Data Fig. 9e,f). Furthermore, a simple statistical model suggests that low dimensionality, long-range correlations and pronounced fractures are intimately connected (Extended Data Fig. 10). Moreover, the effect of heterogeneity in the lateral connections appears to be independent of the specific form of network interactions generating modular activity (Extended Data Fig. 11), and as such, we expect other forms of heterogeneity in connectivity or cellular properties to have a similar effect. Notably, the model predicts that the spatial structure of the correlation peak around the seed point is anisotropic, which is confirmed quantitatively by our data (Fig. 4l; Extended Data Fig. 9g,h). Thus, our dynamical model describes a plausible mechanism for how the early cortex, even in the absence of long-range horizontal connections, could produce spontaneous activity that is correlated over large distances.

Together, our results demonstrate that early cortical circuits generate structured long-range correlations that are refined over development to form mature distributed functional networks, thereby linking the fine-scale functional architecture with distant network organization. Our dynamical model shows that long-range correlations can arise in the early cortex as an emergent property via multi-synaptic short-range interactions that tend to favor certain spatially extended activity patterns at the expense of others. By confining the space of possible large-scale activity patterns, long-range order is established in the form of distributed coactive domains, explaining our observation of long-range spontaneous correlations in the early visual cortex. These correlations appear to possess the required spatial organization, intensity and frequency to guide the emergence of clustered long-range horizontal connections via a Hebbian plasticity mechanism.

## Acknowledgements

We would like to thank D. Ouimet and V. Hoke for technical and surgical assistance, P. Hülsdunk for assistance registering and motion-correcting imaging data, as well as members of the Fitzpatrick and Kaschube laboratories for helpful discussions. This research was supported by US National Institutes of Health grants EY011488 and EY026273 (D.F.), Bernstein Focus Neurotechnology grant 01GQ0840, (M.K.), BMBF project D-USA-Verbund: SpontVision, FKZ 01GQ1507 (M.K.), the International Max Planck Research School for Neural Circuits in Frankfurt, as well as the Max Planck Florida Institute for Neuroscience.

## Author contributions

All authors designed the study, analyzed the results, and wrote the paper. G.B.S. and D.E.W. performed the wide-field and 2-photon calcium imaging, B.H. and M.K. performed the computational modelling. G.B.S., B.H., and D.E.W. contributed equally to this work.

## Author information

The authors declare no competing financial interests. Correspondence and requests for materials should be addressed to D.F. or M.K.

## Methods

### Animals

All experimental procedures were approved by the Max Planck Florida Institute for Neuroscience Institutional Animal Care and Use Committee and were performed in accordance with guidelines from the US National Institutes of Health. 24 female ferret kits were obtained from Marshall Farms and housed with jills on a 16 h light/8 h dark cycle.

### Viral injections

Viral injections were performed as previously described^6,38,39^. Briefly we expressed GCaMP6s^40^ by microinjecting AAV2/1.hSyn.GCaMP6s.WPRE.SV40 (obtained from University of Pennsylvania Vector Core) into the visual cortex approximately 6-10 days prior to imaging experiments. Anesthesia was induced with either ketamine (12.5 mg/kg) or isoflurane (4-5%), and maintained with isoflurane (1–2%). Atropine (0.2mg/kg) and bupivacaine were both administered, and animal temperature was maintained at approximately 37°C with a homeothermic heating blanket. Animals were also mechanically ventilated and both heart rate and end-tidal CO2 were monitored throughout the surgery. Using aseptic surgical technique, skin and muscle overlying visual cortex were reflected and a small burr hole was made with a hand-held drill (Fordom Electric Co.). Approximately 1μL of virus contained in a pulled glass pipette was pressure injected into the cortex at two depths (∼200µm and 400µm below the surface) over 20 minutes using a Nanoject-II (World Precision Instruments). This procedure reliably produced robust and widespread labelling of the visual cortex, with GCaMP6 expression typically extending over an area >3 mm in diameter (Extended Data Fig. 12).

### Cranial window surgery

To allow repeated access to the same imaging field, chronic cranial windows were implanted in each animal 0-2 days prior to the first imaging session. Animals were anesthetized and prepared for surgery as described above. Using aseptic surgical technique, skin and muscle overlying visual cortex were reflected and a custom-designed metal headplate was implanted over the injected region with MetaBond (Parkell Inc.). Then both a craniotomy (∼5mm) and a subsequent durotomy were performed, and the underlying brain stabilized with a 1.4 mm thick 3 mm diameter stacked glass coverslip^39^. The headplate was hermetically sealed with a stainless steel retaining ring (5/16” internal retaining ring, McMaster-Carr) and glue (VetBond, 3M). Unless the animal was immediately imaged after a cranial window surgery, the imaging headplate was filled with a silicone polymer (Kwik-kast, World Precision Instruments) to protect it between imaging experiments.

### Wide-field epifluorescence and two-photon imaging

Wide-field epifluoresence imaging was achieved with a Zyla 5.5 sCMOS camera (Andor) controlled by μManager^41^. Images were acquired at 15Hz with 4 x 4 binning to yield 640 x 540 pixels. Two-photon imaging was performed with a B-Scope microscope (ThorLabs) driven by a Mai-Tai DeepSee laser (Spectra Physics) at 910 nm. The B-Scope microscope was controlled by ScanImage (Vidreo Technologies) in a resonant-galvo configuration with single-plane images (512 × 512 pixels) being collected at 30 Hz.

In animals imaged after eye opening, phenylephrine (1.25-5%) and tropicamide (0.5%) were applied to the eyes to retract the nictitating membrane and dilate the pupil, and the cornea was protected with regular application of eye drops (Systane Ultra, Alcon Laboratories). The silicon polymer plug overlying the sealed imaging chamber was then gently peeled off. Whenever the imaging quality of the chronic cranial window was found to be suboptimal for imaging, the chamber was opened under aseptic conditions, regrown tissue or neomembrane was removed and a new coverslip was inserted. In some cases, prior to imaging, animals were paralyzed with either vecuronium or pancuronium bromide (0.2 mg/kg/h in lactated Ringer’s, delivered IV).

For imaging experiments in awake animals, animals were habituated to head fixation beginning at least 2 days before imaging. Habituation consisted of exposure to the fixation apparatus for brief periods after which animals were returned to their home cage. For imaging, animals were head fixed and wide-field and two-photon imaging was performed as above. In experiments where both awake and anesthetized imaging were performed, awake imaging was always performed first, followed by anesthesia induction as described above. Awake recordings of spontaneous activity were performed in a darkened room and eye position not monitored.

For anesthetized, longitudinal imaging experiments, anesthesia was induced with either ketamine (12.5 mg/kg) or isoflurane (4-5%), and atropine (0.2mg/kg) was administered. Animals were intubated and ventilated, and an IV catheter was placed in the cephalic vein. In some imaging sessions, it was not possible to catheterize the cephalic vein; in these cases, an IP catheter was inserted. Anesthesia was then maintained with isoflurane (0.5–0.75%).

Following imaging, animals were recovered from anesthesia and returned to their home cages. During recovery, neostigmine was occasionally administered to animals that were paralyzed (0.01-0.1µL/kg per dose).

### Visual stimulation

Visual stimuli were delivered on a LCD screen placed approximately 25–30cm in front of the eyes using PsychoPy^42^. For evoking orientation responses, stimuli were full-field sinusoidal gratings at 100% contrast, at 0.015–0.06 cycles per degree, drifting at 1 or 4 Hz, and presented at each of eight directions of motion, for 5s, repeated 8-16 times. In addition, “blank” stimuli of 0% contrast were also presented. Stimuli were randomly interleaved and were presented for 5s followed by a 5-10s gray screen. Spontaneous activity was recorded in a darkened room, with the visual stimulus set to a black screen.

### Analysis software

Data analysis was performed in Python, ImageJ, and Matlab (The Mathworks).

### Signal extraction for wide-field epifluorescence imaging

To correct for mild brain movement during imaging (especially in the awake state), we registered each imaging frame by maximizing phase correlation to a common reference frame. Furthermore, all imaging experiments acquired during a single day were registered into one reference frame. The ROI was manually drawn around the cortical region with high and robust visually evoked activity. The baseline F0 for each pixel was obtained by applying a rank-order filter to the raw fluorescence trace with the rank between 15 to 70 and the time window between 10 and 30s (values chosen for each imaging session individually, depending on the strength of spontaneous activity). The rank and time window were chosen such that the baseline followed faithfully the slow trend of the fluorescence activity. The baseline corrected spontaneous activity was calculated as *(F-F0)/F0* = ΔF/F0.

The baseline for each pixel for the visually evoked activity was obtained by taking the averaged last 1s of the inter-stimulus interval immediately before stimulus onset. The grating evoked response was then calculated as being the average of the fluorescence δF/F0 over the full stimulus period (5s).

### Event detection

To detect spontaneously active events, we first determined active pixels on each frame using a pixel-wise threshold set to 4-5 standard deviations above each pixel’s mean value across time. Active pixels not part of a contiguous active region of at least 0.01mm^2^ were considered ‘inactive’ for the purpose of event detection. Active frames were taken as frames with a spatially extended pattern of activity (>80% of pixels were active). Consecutive active frames were combined into a single event starting with the first high activity frame and then either ending with the last high activity frame or, if present, an activity frame defining a local minimum in the fluorescence activity. In order to assess the spatial pattern of an event, we extracted the maximally active frame for each event, defined as the frame with the highest activity averaged across the ROI. Importantly, calculating the spontaneous correlation patterns (see below) over all frames of all events preserves their spatial structure (Extended Data Fig. 13).

Imaging sessions in which less than 10 spontaneous events were detected were excluded from further analysis. This threshold was chosen based on randomly sampling (with replacement) a varied number of activity patterns, which revealed that spontaneous correlation patterns (see below) for subsamples of >10 events were highly similar (second-order correlation >=0.5) to those obtained from all events (Extended Data Fig. 14).

## Spontaneous correlation patterns

To assess the spatial correlation structure of spontaneous activity, we applied a Gaussian spatial highpass filter (with SD of Gaussian filter kernel *s*_high_=195µm) to the maximally active frame in each event and down-sampled it to 160 x 135 pixels. The resulting patterns, named spontaneous patterns *A* in the following, were used to compute the spontaneous correlation patterns as the pairwise Pearson’s correlation between all locations ***x*** within the ROI and the seed point *s*

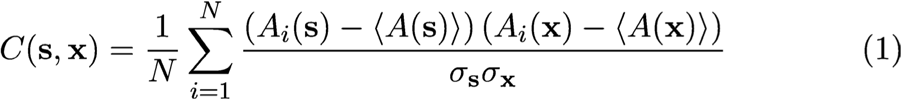

Here the brackets < > denote the average over all events and *σ***_*x*_** denotes the standard deviation of *A* over all *N* events *i* at location ***x***. Note that the spatial structure of spontaneous activity was already evident without filtering (Extended Data Fig. 15). High-pass filtering allowed us to extract this spatial structure, but our results did not sensitively depend on the filtering. For instance, weaker high-pass filtering using a kernel with *s*_high_=520µm resulted in a highly similar correlation structure (data not shown).

### Shuffled control ensemble and surrogate correlation patterns

We compared the real ensemble of spontaneous activity patterns from a given experiment with a control ensemble, obtained by eliminating most of the spatial relationship between the patterns. To this end, all activity patterns were randomly rotated (rotation angle drawn from a uniform distribution between 0° and 360° with a step size of 10°), translated (shifts drawn from a uniform distribution between ±450µm in increments of 26µm, independently for x- and y-direction) and reflected (with probability 0.5, independently at the x- and y-axis at the center of the ROI), resulting in an equally large control ensemble with similar statistical properties, but little systematic interrelation between patterns. Surrogate correlation patterns were then computed from these ensembles as described above.

### Spatial range of correlations

To assess the spatial range of spontaneous correlations (Figs. 1e and 3d), we identified the local maxima (minimum separation between maxima 800µm) in the correlation pattern for each seed point and fitted an exponential decay function

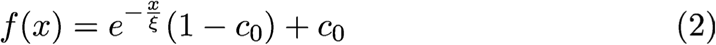

to the values of these maxima as a function of distance *x* to the seed point (Fig. 1e; Extended Data Fig 9a). Here ξ is the decay constant, named ‘spatial scale correlation’ in Fig. 3d and 4j-l. The baseline *c*_0_ accounts for spurious correlations due to a finite number of spontaneous patterns and was estimated as the average value at maxima in the surrogate correlation patterns described above.

To assess the statistical significance of long-range correlations ∼2 mm from the seed point, we compared the median correlation strength for maxima located 1.8-2.2 mm away against a distribution obtained from 100 surrogate correlation patterns. For individual animals, the p-value was taken as the fraction of median correlation strength values from surrogate data greater than or equal to the median correlation strength for real correlation patterns. For 2 of 12 animals, the statistical significance of long-range correlations could not be assessed, due to insufficient coverage in rotated and translated surrogate activity patterns caused by an irregularly shaped ROI. These animals were excluded from analysis of long-range correlation strength.

### Comparison of awake and anesthetized correlations

Correlation similarity across awake and anesthetized states was computed for each seed-point as the Pearson’s correlation coefficient of the spontaneous correlations for that seed point across states. For each seed-point, correlations within 400µm were excluded from analysis. These “second-order correlations” (shown for each seed point in Extended Data Fig. 4f) were then averaged across all seed points within the ROI. To determine the significance of these second-order correlations across state, we shuffled corresponding seed points across states 1000 times, and again computed correlation similarity. Likewise, to gain an estimate of the expected similarity for a well-matched correlation structure, we computed the similarity of each state to itself. Correlation patterns were first separately computed for half of the detected events, and then the two patterns were compared as above.

### Comparison of wide-field and cellular correlations

2-photon images were corrected for in plane motion via a 2D cross correlation-based approach. For awake imaging, periods of excessive motion were discarded and excluded from further analysis. Cellular regions of interest (ROIs) were drawn using custom software (Cell Magic Wand, ^43^) in ImageJ and imported into Matlab via MIJ ^44^. Fluorescence was averaged over all pixels in the ROI and traces were converted to δF/F0 ^6^, where the baseline fluorescence, F0, was computed from a filtered fluorescence trace. The raw fluorescence trace was filtered by applying a 60s median filter, followed by a first-order Butterworth high-pass filter with a cut-off time of 60s.

To compute spontaneous correlations (Fig. 1f, g), we first identified frames containing spontaneous events, which were defined as frames in which > 30% of imaged neurons exhibited activity > 2 standard deviations above their mean. The stability of activity during an event was computed as the cross-correlation of each frame with the peak activity frame, and was compared to a distribution of 100 randomly chosen intervals of the same length. Cellular activity on all event frames was then Z-scored using the mean and standard deviation of each frame, and correlation patterns for each cell were computed as the pairwise Pearson’s correlation coefficient, using the activity of all neurons on all active frames.

To compare the correlation structure obtained at the cellular level with that obtained via wide-field imaging (Fig. 1g) we first aligned the 2-photon field of view (FOV) to the wide-field image using blood vessel landmarks and applied an affine transformation to obtain the pixel coordinates of each imaged neuron in the wide-field frame of reference. Correlation similarity was obtained as above by computing the second-order correlation between the cellular correlation structure and that of the corresponding wide-field pixels, using all cells >200µm from the seed point. Shuffled second-order correlations were obtained by randomly rotating and translating the 2P FOV within the full wide-field ROI, 1000 times. To estimate the maximum expected degree of similarity, we computed a second-order correlation within the cellular correlation structure itself by determining the similarity of correlation structures computed using only 50% of detected events (dashed line and blue bar in Fig. 1i).

### Orientation preference and ocular dominance maps

The orientation preference maps (Fig. 2a, 3f (*right*)) were calculated based on the trial-averaged responses evoked by binocularly presented moving grating stimuli of eight directions equally spaced between 0 and 360 degree. Responses were Gaussian band-pass filtered (SD: *s*_*low*_=26µm, *s*_*high*_=195µm) and orientation preference was computed by vector summation:

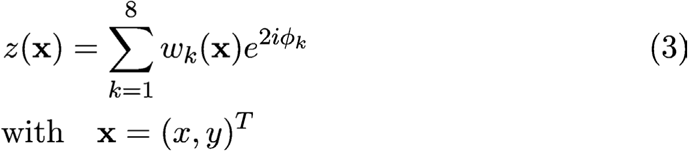

where *w*_*k*_(***x***) is the tuning curve at location ***x***, i.e. the trial-averaged response to a moving grating with direction ⁏_*k*_ at location ***x***. The preferred orientation at ***x*** is 0.5 arg*(z(****x***)).

Orientation pinwheel centers (Fig 2i, j) were estimated as described in Refs. ^45,46^. The Matlab routine provided by Schottdorf et al. ^46^. was used. Orientation contour lines (Fig. 2b, 3f) are the zero-levels of the 0°-90° difference map, obtained by using the matplotlib.pyplot.contours routine. Surrogate orientation preference maps were obtained by phase shuffling the original maps in the Fourier domain^45^.

Ocular dominance maps were calculated based on the trial-averaged responses evoked by presenting moving grating stimuli of eight directions equally spaced between 0 and 360 degree either to the contralateral or ipsilateral eye. The trial averaged response to each orientation and ocular condition was Gaussian band-pass filtered as described above for the orientation map. Contralateral and ipsilateral response maps were computed by respectively averaging together the trial-average responses to the stimuli presented either to the contralateral or ipsilateral eye. The ocular dominance map was computed as a difference of the contralateral and ipsilateral response maps.

### Similarity of correlation patterns to the orientation and ocular dominance maps

To quantify how similar patterns of correlated spontaneous activity are to known functional maps in visual cortex, we computed the average pairwise similarity of the spontaneous correlation patterns either to the ocular dominance map or the orientation preference map (Extended Data Fig. 5). The assessment of similarity of each correlation pattern to the ocular dominance map is the magnitude of their pairwise coefficient:

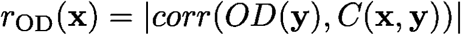

where *OD(****y***) is the ocular dominance map at location ***y*** and *C(****x, y***) is the spontaneous correlation pattern between seed location ***x*** and location ***y*** and *corr* denotes Pearson’s correlation coefficient. Correspondingly, the similarity of each correlation pattern to the orientation map is computed as the magnitude of the pairwise correlation coefficient to the real and imaginary components of the vector-summed orientation map *z*:

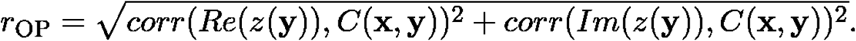

### Prediction analysis and exclusion areas

To test whether orientation tuning can be predicted from the tuning at remote locations with correlated spontaneous activity (Fig. 2c), we estimated the tuning curve at seed point ***s****=(s*_*x*_,*s*_*y*_) by the sum over tuning curves *w*_*k*_ at different locations weighted by their spontaneous correlation *C* with the seed point:

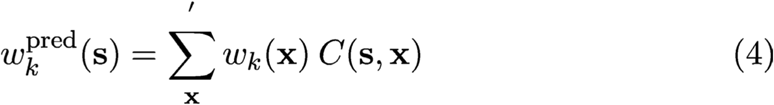

where *k* denotes the orientation of the stimulus. The sum was taken over locations ***x*** outside a circular area centered at the seed point with radius 0.4, 1.2 or 2.4mm. For this calculation, both *w*_*k*_ and *C* are z-scored. To assess the goodness of the prediction, we calculated the angular difference between the predicted and the actual preferred orientation (Fig. 2f). Low values indicate a high match, whereas 45° indicates chance level.

Statistical significance (Fig. 2f) was determined by repeating this analysis for 100 surrogate orientation preference maps, obtained by phase shuffling in the Fourier domain^45^. For individual animals, the p-value was taken as the fraction of values equal or smaller than the value for the real orientation map. To pool across animals within an exclusion radius (Fig. 2f), we then generated 10,000 surrogate group medians by randomly drawing from the distributions of surrogate data points (one per animal), and the p-value was taken as the fraction of group medians equal or smaller than the median value for the actual data.

### Spontaneous fractures

Fracture strength was defined as the rate by which the correlation pattern changes when changing the seed point location over some small distance (Fig. 2h (*bottom*), Fig. 2i). It was computed as:

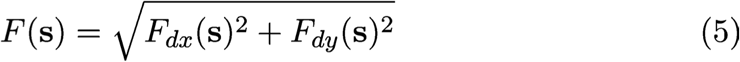

where *F*_*dx*_ (*F*_*dy*_) denotes the x(y)-component of the rate of change of the correlation pattern at seed point ***s***. We approximated this rate of change by the (second-order) correlation between two correlation patterns with seed points at adjacent pixels a distance *d* apart:

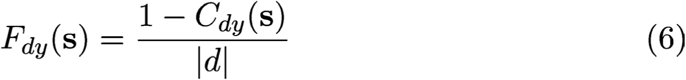

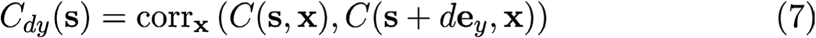

where corr**_*x*_** denotes Pearson’s correlation coefficient calculated over all locations ***x*** and ***e***_*y*_ is a unit vector in y-direction. The subtraction from 1 in the numerator ensures *F=0* at seed point locations, around which the correlation pattern does not change, while high values of *F* indicate high changes. We used *d*=26µm, the spatial resolution of the correlation patterns.

We defined fracture magnitude (Fig. 4e (*bottom*), Fig. 4i) as the difference between *F*, averaged over the fracture lines, and its average in regions *>*130µm apart from the nearest fracture line. To extract the fracture lines from *F* we first applied a spatial median filter with a window size of 78µm to remove outliers. We then applied histogram normalization, contrast enhancement by using Contrast Limited Adaptive Histogram Equalization (CLAHE, clip limit=20, size of neighborhood 260×260µm^2^), and a spatial highpass filter (Gaussian filter, SD *s*_*high*_=390µm). The resulting values were binarized (threshold=0), and the resulting two-dimensional binary array eroded and then dilated (twice) to remove single not-contiguous pixels. We skeletonized this binary array to obtain the fracture lines.

We quantified the co-alignment between spontaneous fractures and high orientation gradient regions by the fracture selectivity (Fig. 2k), defined as the difference between *F* at high orientation gradient locations (***x***_*high*_, >π/5 radians/pixel) and locations far from high orientation gradients (***x***_*low*_, >150µm from ***x***_*high*_):

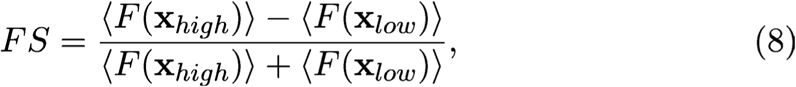

where the brackets denote average over locations ***x***_*high*_ and low ***x***_*low*_, respectively. A value of *FS* of 1 indicates co-alignment between the spontaneous fractures and the orientation gradient, whereas a value near 0 indicates no such alignment. To assess significance (Fig. 2k) we repeated this analysis for 1000 surrogate orientation preference maps, obtained by phase shuffling in the Fourier domain. The p-value is the fraction of values equal or larger than the value for the orientation map.

In order to test whether spontaneous fractures reflect the correlation structure over remote distances and not only in their local neighborhood (Fig. 2i (*top*), Fig. 2l), we computed *F* as above, but excluding a circular region with radius 0.4, 1.2 or 2.4 mm, centered at the seed point ***s***. We then computed the Pearson’s correlation coefficient with the original *F*.

### Registration for longitudinal imaging

To compare spontaneous correlation patterns across days in longitudinally imaged animals, we transformed all imaging data into a common reference frame (Extended Data Fig. 6a). This transformation corrected for small displacement and expansion of cortical tissue over the imaging period, presumably due to cortical growth. We used an affine transformation, thereby taking into account rotation, scaling, translation and shear mapping of the cortex:

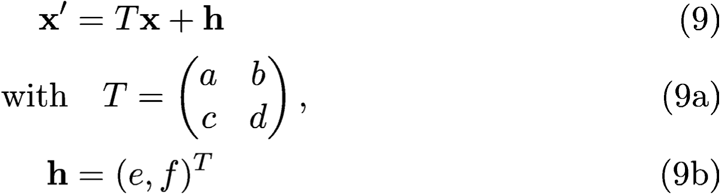

The parameter of the transformation matrix *T* and of the displacement vector ***h*** were found by minimizing the distance between landmarks determined for each day of experiment. Landmarks were found by marking radial blood vessels (i.e. blood vessels oriented orthogonally to the imaging plane) by visual inspection. The following expression was minimized (least square fit) to find transformation parameters from day *t* to the reference day *t*_*ref*_ (eye opening):

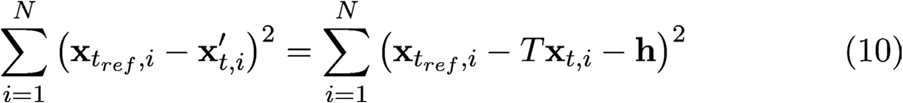

with *N* landmarks (between 10 to 30) in both coordinate systems at coordinates ***x***_*tref,i*_ in the reference coordinate system, and the coordinates ***x***_*t,i*_ at day *t*.

### Analysis of spontaneous correlation across development

To compare spontaneous correlation patterns across development, we calculated a second-order correlation (Extended Data Fig. 6d,e) between the correlation patterns on a given day and the reference day (eye opening) with the same seed point. Changes in correlation fractures over development were quantified as the second order correlation of fracture patterns (Extended Data Fig. 6f). In both cases, an estimate of the expected degree of similarity was computed by first separately computing correlations and their corresponding fracture patterns for half of the detected events, and then computing the second-order correlations as above.

To determine whether correlation patterns early in development can predict mature orientation preferences (Fig. 3h), we computed orientation tuning predictions as above, using the correlation pattern on a given day to weight tuning curves measured following eye opening, with an exclusion radius of 400 µm. The predicted orientation preference map was compared to the actual map as described above (“Prediction analysis and exclusion areas”) for both individual animals and group medians.

To assess the statistical significance of long-range correlation strength at 2 mm across development, we compared correlation maxima to those of surrogate correlation patterns as described above (“Spatial range of correlations”). To pool across experiments within an age group, we then generated 10,000 surrogate group medians by randomly drawing from the distributions of surrogate data points (one per experiment), and the p-value was taken as the fraction of group medians greater than the median value for the actual data.

### Retinal and LGN inactivation experiments

For retinal inactivation experiments, a cranial window was implanted over visual cortex as described above. After imaging spontaneous activity under light isoflurane anesthesia (0.5-1%) as described above, visually evoked responses were recorded in response to full-field luminance steps^6^. Isoflurane levels were then increased and intraocular infusions of TTX were performed into each eye. For each intraocular injection, a small incision was made just posterior to the scleral margin using the tip of a 30-gauge needle attached to a Hamilton syringe. Each eye was then injected with 2-2.5 µL of 0.75 mM TTX solution (Tocris Bioscience) to reach an intraocular dose of 21.45 µM that is roughly comparable the dosage used previously in the ferret^47^. Following infusion of TTX, isoflurane levels were reduced, and the animal returned to a stable light anesthetic plane. The efficacy of TTX was tested by the absence of visually evoked responses to full-field luminance steps. Following confirmation of retinal blockade, spontaneous activity was imaged as above. Following collection of spontaneous activity, retinal blockade was again confirmed through the absence of cortical responses to visual stimuli.

For LGN inactivation experiments, surgical preparation was as described above. A head-post was implanted near bregma, a craniotomy was made over visual cortex, and sealed with a coverslip affixed directly to the skull with cyanoacrylate glue and dental cement. A second craniotomy was then made over the approximate location of the LGN (Horsley-Clarke coordinates: AP -1mm, LM 6mm). The LGN was typically located at a depth of 5-8.5mm, and its spatial position mapped by identifying units responsive to a full-field luminance stimulus through systematic electrode penetrations. Once the LGN position was determined, spontaneous activity in visual cortex was recorded as above, followed by visually-evoked responses to luminance steps. A micropipette filled with muscimol (25-100 mM, Tocris Biosciences) was lowered into the center of the LGN, and infusions of ∼0.5 µL were made at three depths along the dorsal-ventral extent of the penetration using a nanoliter injector (Nanoject). The efficacy of thalamic inactivation was confirmed by the abolishment of visually evoked activity prior to and following imaging of spontaneous activity in the cortex.

Spontaneous activity was analyzed as described above, with one exception: the 10 event threshold for inclusion (see above) was not applied to the LGN inactivation experiments as in 1 of 3 cases <10 events were recorded following LGN inactivation. Comparisons between pre- and post-inactivation patterns made using second-order correlations as described above for comparisons of awake and anesthetized activity.

### Local correlation structure

To describe the shape of the peak of a correlation pattern around its seed point (Fig. 4l; Extended Data Fig. 9g), we fitted an ellipse (least-square fit) with orientation *φ*, major axis *ς*_*1*_ and minor axis *ς*_*2*_ to the contour line at correlation=0.7 around the seed point. The eccentricity *ε* of the ellipse is defined as:

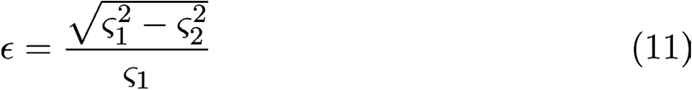

Its value is 0 for a circle, with increasing values indicating greater elongation of the ellipse.

#### Dimensionality of spontaneous activity

We estimated the dimensionality *d*_eff_ of the subspace spanned by spontaneous activity patterns (Fig. 4a) by (see Ref ^48^):

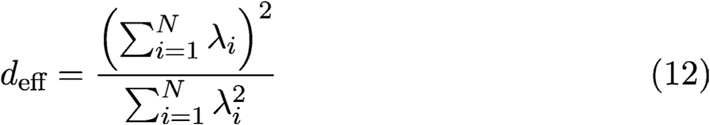

where *λ*_*i*_ are the eigenvalues of the covariance matrix for the N locations (pixels) within the ROI (Extended Data Fig. 9e). These values were compared to the dimensionality of surrogate spontaneous activity patterns by taking the median value of 100 surrogate ensembles generated for each animal as described above (in “Shuffled control ensemble and surrogate correlation patterns”).

#### Statistical Model

To generate a statistical ensemble of spatially extended, modular patterns with predefined dimensionality *k* (Extended Data Fig. 10), we first synthesized *k* two-dimensional Gaussian random fields^45^ with spectral width matched to that of the experimentally observed spontaneous activity patterns (size 100×100 pixel; spatial period *Λ*_*stat*_ 10 pixel). Interpreting these *k* patterns as vectors ***v***_*j*_ (*j*=1,…,k) in the high-dimensional pixel space, we orthonormalized them based on a Householder reflection. From these *k* orthonormal basis vectors ***v***, we generated activity patterns *A*_*i*_ (*i*=1, … 10,000) by linear combinations with coefficients ζ drawn independently from a Gaussian distribution with 0 mean and SD equal to 1:

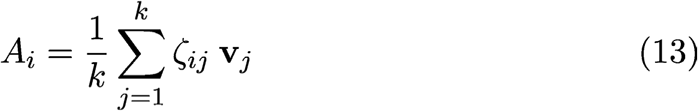

Over this ensemble we computed the correlation patterns analogously to the analysis of the experimental data. From these we computed the fracture strength and magnitude (Extended Data Fig. 10f, (*bottom*)) and the local maxima at a distance of 4*Λ*_*stat*_ from the seed point to estimate the strength of long-range correlations (Extended Data Fig. 10f, (*top*)).

#### Dynamical Model

The model addresses the question whether short-range lateral connections can give rise to patterns of spontaneous activity that are (i) modular, exhibit (ii) long-range correlations and (iii) pronounced spontaneous fractures. Feature (i) can be explained by the Turing-mechanism, which for simplicity we implemented employing a ‘Mexican hat’ connectivity (local excitation, lateral inhibition), but other network motives can be used as well (Extended Data Supplementary Notes). The model shows that heterogeneity in the lateral connections is sufficient to explain features (ii) and (iii). Heterogeneity was implemented using elongated Mexican hats whose properties vary randomly across cortex.

We modeled the early spontaneous activity by a two-dimensional firing rate network (Fig. 4i-l) obeying the following dynamics

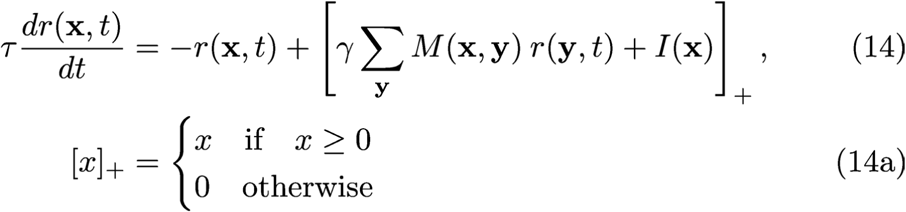

where *r(****x***,*t)* is the average firing rate in a local pool of neurons at location ***x***, *τ* is the neuronal time constant, *M(****x,y***) are the synaptic weights connecting locations ***x*** and ***y,*** *I(****x***) is the input to location ***x***, and γ a factor controlling the overall strength of synaptic weights. The connectivity *M* is assumed to be short-range and follows a Mexican hat structure. Moreover, the Mexican hats are anisotropic, modeled as the difference of two elongated Gaussians, whose axis of elongation and scale vary discontinuously across space:

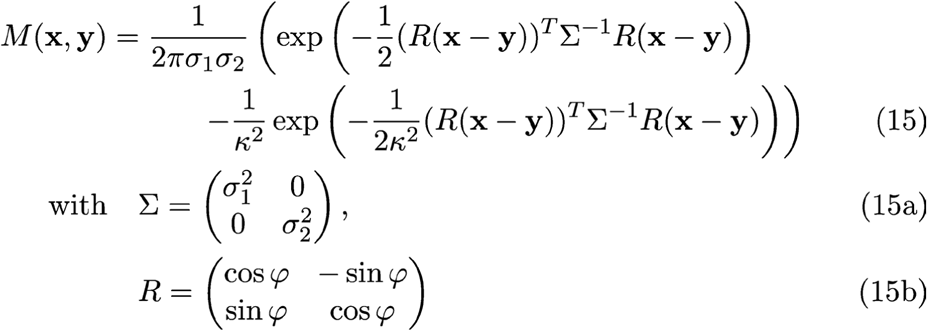

Here *σ*_*1*_ and *σ*_*2*_ (≤ *σ*_*1*_) denote the SDs of the smaller Gaussian in the direction of its major and minor axis, respectively. For the larger Gaussian both SDs are scaled by a factor *κ≥1*. The level sets of the Gaussians are ellipses whose larger (smaller) axis is proportional to *σ*_*1*_ (*σ*_*2*_) and so the eccentricity *ε* (eq. 11) measures the degree of elongation of the Mexican hat. The angle *φ* determines the orientation of the elongated Mexican hat. The dependence of these parameters on cortical space **x** is suppressed in eq. 15 for brevity. *M* is normalized such that the magnitude of its maximal eigenvalue is equal to 1. For all simulations we set *κ*=2, *τ*=1 and γ=1.02 and used random initial conditions *r(****x***,*t=0)* drawn from a uniform distribution between 0 and 0.1.

We introduced the heterogeneity parameter *H* to parameterize and systematically vary the heterogeneity of elongated Mexican hats across cortical space **x**. The eccentricity *ε* was drawn from a normal distribution with mean <*ε*> and standard deviation *σ*_*ε*_ both depending linearly on *H* (<*ε>=H, σ_ε_=*0.13 *H*). The size of *σ*_*1*_ was drawn from a normal distribution with SD 0.1<*σ*_*1*_>*H* and mean <*σ*_*1*_>=1.8. The orientation *φ* of the Mexican hat axis was drawn from a uniform distribution between 0° and 180°. These three parameters were drawn independently at each location.

In the case of isotropic Mexican hat connectivity (*σ*_*1*_=*σ*_*2*_) the eigenvectors of *M* are plane waves and the spectrum is peaked at the wavenumber *k*=2π/*Λ*, and thus the typical spatial scale *Λ* of the pattern is given by

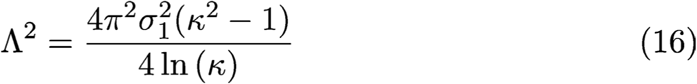

This defines the spatial scale *Λ* used as reference in Fig. 4i,j. For comparison between model and data we identified 1*Λ* with 1mm, which is roughly the spatial scale of spontaneous patterns observed in experiment.

The input drive *I* is assumed constant in time for simplicity, consistent with our observation that in the early cortex spontaneous patterns were often fairly static during a spontaneous event (Extended Data Figure 2 and Movie 5). *I* is modulated in space using a band-pass filtered Gaussian random field *G* with spatial scale *Λ,* zero mean and unit SD^45^:

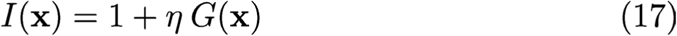

We varied the input modulation *η* between 0.004 and 0.4 in Figs. 4k,l the regime over which we observed a smooth transition from an input-dominated system to a system dominated by the recurrent connections.

To model a spontaneous event, we integrated eq. 14 until a near steady state of the dynamics was reached. The results in Fig. 4j-l were obtained for an integration time of 500*τ*, but already a much shorter integration over 50*τ* resulted in similar solutions and nearly the same level of long-range correlations and dimensionality. Different spontaneous events were obtained by using different realizations of input drive *I* and initial conditions (same connectivity *M*). To generate Figs. 4j(bottom),k,l and Extended Data Figs. 9b,d,f,h and 11 we furthermore averaged over 10 realizations of connectivity *M* for each parameter setting.

We numerically integrated the dynamics using a 4^th^ order Runge-Kutta method in a square region of size 100 x 100 using periodic boundary conditions. The time step was *dt*=0.15*τ* and the spatial resolution 10 pixel per *Λ*.

The simulations were performed on the GPUs GeForce GTX TITAN Black and GeForce GTX TITAN X. The code was implemented in Python and Theano (version 0.8.1).

#### Statistical analysis

Non-parametric statistical analyses were used throughout the study. All tests were two-sided unless otherwise noted. Bootstrapping and surrogate approaches were used to estimate null distributions for test statistics as described above. Sample sizes were chosen to be similar to prior studies using similar methodologies in non-murine species (e.g. Refs: ^6,12,13,26,38^). All animals in each experiment were treated equivalently, and no randomization or blinding was performed.

#### Code availability

The code for data analysis and simulations can be made available upon request to the corresponding author.

#### Data availability

The data that support the findings of this study are available from the corresponding author upon reasonable request.

